# Development of a mouse model for spontaneous oral squamous cell carcinoma in Fanconi anemia

**DOI:** 10.1101/2022.06.05.494848

**Authors:** Ricardo Errazquin, Angustias Page, Anna Suñol, Carmen Segrelles, Estela Carrasco, Jorge Peral, Alicia Garrido-Aranda, Sonia Del Marro, Jessica Ortiz, Corina Lorz, Jordi Minguillon, Jordi Surralles, Cristina Belendez, Martina Alvarez, Judith Balmaña, Ana Bravo, Angel Ramirez, Ramon Garcia-Escudero

**Affiliations:** Research Institute Hospital 12 de Octubre (imas12), University Hospital “12 de Octubre”, Av Córdoba s/n, 28041 Madrid, Spain; Centro de Investigación Biomédica en Red de Cáncer (CIBERONC), 28029 Madrid, Spain; Biomedical Oncology Unit, CIEMAT (Centro de Investigaciones Energéticas, Medioambientales y Tecnológicas), Avenida Complutense 40, 28040 Madrid, Spain; Hereditary Cancer Genetics Group and Medical Oncology Department, VHIO, Barcelona, Spain; Centro de Investigaciones Médico-Sanitarias (CIMES), Malaga, Spain; Join Research Unit on Genomic Medicine UAB-Sant Pau Biomedical Research Institute (IIB Sant Pau), Hospital de la Santa Creu i Sant Pau, 08041 Barcelona, Spain; Centro de Investigación Biomédica en Enfermedades Raras (CIBERER), 28029 Madrid, Spain; Pediatric Hematology and Oncology. Hospital General Universitario Gregorio Marañón, Madrid, Spain; Facultad de Medicina, Universidad Complutense de Madrid, Spain; Instituto Investigación Sanitaria Gregorio Marañón, Madrid, Spain; Department of Anatomy, Animal Production and Veterinary Clinical Sciences, Laboratory of Pathology Phenotyping of Genetically Engineered Mice, Faculty of Veterinary Medicine, University of Santiago de Compostela, 27002 Lugo, Spain

**Keywords:** Fanconi anemia, head and neck squamous cell carcinoma, oral squamous cell carcinoma, mouse model, oral mucosa, FANCA, TP53, Trp53, p53, mutation

## Abstract

Fanconi anemia (FA) patients frequently develop oral squamous cell carcinoma (OSCC). This cancer in FA patients is diagnosed within the first 3-4 decades of life, very often preceded by lesions that suffer a malignant transformation. In addition, they respond poorly to current treatments due to toxicity or multiple recurrences.

Translational research of new chemopreventive agents and therapeutic strategies has been unsuccessful partly due to scarcity of disease models or failure to fully reproduce the disease. Here we report that Fanca gene knockout mice (Fanca-/-) frequently display pre-malignant lesions in the oral cavity. Moreover, when these animals were crossed with animals having conditional deletion of *Trp53* gene in oral mucosa (*K14cre*;*Trp53*^F2-10/F2-10^), they spontaneously developed OSCC with a high penetrance and a median latency of less than ten months. Tumors were well differentiated and expressed markers of squamous differentiation, such as keratins K5 and K10. In conclusion, *Fanca* and *Trp53* genes cooperate to suppress oral cancer in mice, and *Fanca*^*-/-*^*;K14cre;Trp53*^F2-10/F2-10^ mice constitute the first animal model of spontaneous OSCC in FA.

## INTRODUCTION

Fanconi anemia (FA) is a heritable syndrome with predisposition to congenital abnormalities, bone marrow failure (BMF), and cancer. In FA patients, pathogenic mutations have been described in 23 different FA genes which give rise to Fanconi or Fanconi-like clinical phenotype. Most FA children have BMF that can be successfully treated with hematopoetic stell cell transplant (HSCT). HSCT has also been used to treat blood malignancies such as myelodysplastic syndrome (MDS) and leukemia in FA. HSCT has significantly improved life expectancy in FA individuals so that there are now more adults living with FA than children diagnosed with this syndrome. Unfortunately, a consequence of this achievement has been the identification of solid tumor predisposition as the most important health challenge in FA young adults. Patients have an extraordinarily high lifetime risk of squamous cell carcinomas (SCC) of the head and neck, esophagus, vulva, or anus. This risk is increased by prior HSCT, together with a continuing risk of acute myeloid leukemia [1]. Synchronic and metachronic tumors, especially in the oral cavity, are distressingly common in FA [2]. Besides SCC, other malignant tumors have also been described in FA, although less frequently: lymphoma, liver cancer, brain tumors, and embryonal tumors such as Wilms tumor and neuroblastoma. However, the risk of these types of tumors is much lower than SCC.

A hallmark feature of FA is the inability to repair DNA interstrand cross-links (ICLs). FA proteins repair ICLs in a common cellular pathway known as the FA pathway or FA/BRCA pathway. A defective FA pathway leads to the accumulation of genomic aberrations and the emergence of precancerous cells that eventually might progress into invasive carcinomas. However, the etiology of SCC in mucosal epithelia in FA is not well understood. Known drivers of head and neck SCC (HNSCC) are smoking and alcohol consumption, but they are less commonly reported in FA than in the general population. Human papillomavirus (HPV) infection is another driver of HNSCC. However, some reports suggest that HPV may be a major contributor to HNSCC development in patients with FA [3], whereas other studies dispute these results [4]. HNSCC in FA is difficult to treat, as patients cannot tolerate current therapies, including ionizing radiation and platinum-based chemotherapies [5]. In addition, most tumors are detected at advanced stages. Altogether, these events have made cancer the principal cause of early mortality in adults with FA. Therefore, the discovery of non-genotoxic and efficient treatment opportunities for FA patients is of utmost importance.

New therapies against FA HNSCC could be discovered and preclinically tested in adequate disease models such as cellular and animal models. Tremendous efforts have been made to generate and analyze whether genetically engineered mouse models (GEMMs) with deleterious mutations in FANC genes could model disease features of FA patients. Thus, lines of mice bearing mutations in *Fanca, Fancc, Fancd1, Fancd2, Fance, Fancf, Fancg, Fanci, Fancl, Fancm, Fancn, Fanco*, or *Fancp* have been produced [6-8]. To date, all FA mouse models display reduced fertility and cells derived from them are sensitive to ICLs. Embryonic and perinatal lethality has been observed in most FA models, which could be influenced by the mouse genetic background. Some models display other minor developmental abnormalities, such as microphthalmia in *Fanca* [9], *Fancc* [10], *Fancd2* [11], *Fanci* [7], and *Fancp* [12]. The life-threatening anemia that FA patients suffer is not modelled in FA mice, although some defects are present, such as lower blood cellularity in Fancp [12] or reduced proliferation in hypomorphic *Fancd1* [13]. Futhermore, the high incidence of BMF, leukemia and SCC, accompanied with early mortality in adults with FA, is not well recapitulated in FA mice. Long-term survival has been reported in *Fancc*-, *Fancd2*-, *Fancf*-, *Fancm*- and *Fancp*-deficient mice. However, increased incidence of various solid tumors exists in *Fanca*-, *Fancf*-, *Fancd2*- and *Fancm*-mice. Strikingly, none of these animals develop SCC in the oral cavity, esophagus or anogenital tissues. These findings demonstrate that the loss of a single *Fanc* gene is not a sufficient condition to model early cancer appearance or SCC development in GEMMs. Double-mutant mice have been investigated to uncover genetic interactions with other cellular pathways leading to oncogenesis. Cooperation in tumor suppression clearly exists between the FA pathway and p53. Accelerated tumorigenesis is present in double *Fancc*^*-/-*^ *;Trp53*^*-/-*^ [14] and *Fancd2*^*-/-*^*;Trp53*^*-/-*^ [15], although these mice do not show increased SCC formation in oral and anogenital tissues, thus precluding their use as animal models of SCC in the appropriate tissue of origin. Fancd2-/- animals, when crossed with *K14cre;HPVE6* or *K14cre;HPVE6E7*, and upon treatment with oral carcinogen 4-NQO, developed HNSCC [16, 17]. In these animals, no tumor formation was reported in the absence of 4-NQO.

Most HNSCCs from FA patients display alterations typically found in HPV-negative HNSCC tumors from the general population, such as mutations in the *TP53* gene [18-21]. Here, we also detected mutations in *TP53* in all five SCCs analyzed from the oral cavity of three FA patients. In order to develop a mouse model of FA oral SCC, we decided to cross *Fanca*^-/-^ animals with *K14cre*;*Trp53*^F2-10/F2-10^ mice lacking p53 in stratified epithelia (such as oral mucosa). We previously reported that K14*cre*;*Trp53*^F2-10/F2-10^ animals spontaneously develop skin SCC, demonstrating a predominant tumor suppressor role of p53 in mouse epidermis [22]. Furthermore, *K14cre*;*Trp53*^F2-10/F2-10^ mice display oral cavity tumors when combined with constitutive Akt activity [23] or IKKβ overexpression [24]. We then hypothesized that combined loss of *Fanca* and *Trp53* genes in the oral cavity could produce spontaneous oral tumors. Here we describe the phenotype of double *Fanca*^-/-^; *K14cre*;*Trp53*^F2-10/F2-10^ mice and we demonstrate that loss of both *Fanca* and *Trp53* genes in oral mucosa recapitulates human FA HNSCC.

## MATERIALS AND METHODS

### Patients

Oral squamous cell carcinomas (OSCCs) from 3 Fanconi anemia patients were analyzed. The study was conducted in accordance with the precepts established in the Declaration of Helsinki, Good Clinical Practice guidelines, and all applicable regulatory requirements. The ethics committee of Hospital 12 Octubre approved the study protocol (study number 18/294).

### Tumor sequencing from FA patients

Deep-sequencing was performed from formalin-fixed paraffin-embedded (FFPE) tumor blocks, from 10-15 sections of 4 µm thickness. Briefly, the cancerous tissue on 4–5 slides (4 µm sections) per patient was dissected out. These were then deparaffinated using deparaffinisation solution (Qiagen, Hilden, Germany), and areas containing >30% tumor cells as determined by an expert pathologist were macrodissected and deparaffinated using Deparaffinization Solution (Qiagen, Hilden, Germany). DNA was extracted and purified with GeneRead DNA FFPE Kit (Qiagen, Hilden, Germany). DNA concentration was quantified using Qubit™ ds DNA High-Sensitive Assay kit (Thermo Fisher Scientific, Waltham, MA, USA) on the Qubit fluorometer (Thermo Fisher Scientific, Waltham, MA, USA). All library preparation was performed manually for Oncomine Comprehensive Assay v3 (OCAv3) (Thermo Fisher Scientific, Waltham, MA, USA) according to manufacturer’s instructions MAN0015885. Multiplex PCR amplification was conducted using a DNA concentration of approximately 20 ng as input. For sequencing, prepared libraries were loaded according to manufacturer’s instructions (Ion 540™—Chef, MAN0010851) onto Ion 540™ Chips (Thermo Fisher Scientific, Waltham, MA, USA) and prepared using the Ion Chef™ System. Sequencing was performed using the Ion GeneStudio™ S5 Prime System (Thermo Fisher Scientific, Waltham, MA, USA). The data was mapped to the human genome assembly 19, embedded as the standard reference genome in the Ion Reporter™ Software (v. 5.14) (Thermo Fisher Scientific, Waltham, MA, USA). We used the Ion Reporter™ Software for initial automated analysis, and Oncomine Comprehensive v3—w4.0—DNA—Single Sample as analysis workflow. Additionally, coverage analysis reports from the Ion Reporter™ Software providing measurements of mapped reads, mean depth, uniformity, and alignment over a target region were used as quality assessment of the sequencing reactions.

### Mice

*Fanca*^-/-^ animals were maintained in FVB/NJ background and obtained from the laboratory of Dr. Paula Rio (Biomedical Innovation Unit, CIEMAT, Madrid, Spain) [25]. *Trp53*^F2-10/F2-10^ and *K14cre* mice were maintained in a mixed FVB/NJ x DBA/2J x C57BL/6 background. Animals were genotyped by PCR using specific primers as previously described [22, 25, 26]. All animals showing tumors, or significant morbidity, over a period of 24 months were sacrificed for necropsy. Histopathological analysis was systematically performed on the esophagus and the oral cavity, including lips, palate, tongue, and oral mucosa. Overt tumors existing in other organs were also sampled for histopathology. All mice husbandry and experimental procedures were performed according to European and Spanish regulations and were approved by the local Animal Ethical Committee and competent authority (code PROEX006/18).

### Histopathology and Immunohistochemistry

Mouse tissues were fixed in 4% buffered formalin and embedded in paraffin. Sections (5 microns) were stained with H&E or processed for immunostaining. Sections were incubated with primary antibodies and thereafter with biotinylated secondary antibodies (Jackson Immunoresearch Laboratory, Ely, UK). Primary antibodies used were: anti-Keratin 5 polyclonal antibody, clone Poly19055 (BioLegend); anti-Keratin 10 antibody, clone Poly19054 (BioLegend). Immunoreactivity was revealed using the ABC-peroxidase system and the DAB substrate kit (Vector Laboratories; Burlingame, CA, USA), and the sections were counterstained with hematoxylin. Anti-Cytokeratin 5 antibody was purchased from Santa Cruz (sc-32721). The control experiments without the primary antibody gave no signal.

### Survival Curves

Tumor-free survival curves were obtained with Prism software (Graphpad Software, Inc., San Diego, CA, www.graphpad.com). Statistical significance of survival between genotypes was calculated with the log-rank test yielding a p-value.

## RESULTS

### Oral squamous carcinomas from Fanconi anemia patients display mutations in *TP53*

We performed deep-sequencing of DNA from cancerous areas of five FFPE blocks from three FA patients who had been diagnosed with squamous cell carcinoma in the oral cavity and treated surgically. Patient clinical and sample information is shown in Supplementary Table 1. Genes included in the sequencing panel are frequently mutated in solid tumors (Supplementary Table 2). Interestingly, mutations in *TP53* previously reported to be oncogenic drivers were detected in all five tumor samples analyzed (Supplementary Table 3 and Supplementary Table 4), as well as mutations and copy number variants in other genes (Supplementary Table 4 and 5). In addition, we also found the germline mutations in the *FANCA* gene that had previously been diagnosed in all three FA patients [27] (Supplementary Table 3). These results are in line with the frequent detection of *TP53* alterations in SCC from FA patients [19-21]. Therefore, animal models resembling germline FANCA and somatic *TP53* mutations might model SCC in FA.

### *Fanca/p53*^EPI^ and *p53*^EPI^ animals develop spontaneous squamous cell carcinomas in the oral cavity (OSCC)

Mice of the indicated genotypes were monitored for a period of 24 months, and spontaneous lesions in the oral cavity were analyzed. We found OSCC in *K14cre*;*Trp53*^F2-10/F2-10^ (hereinafter referred *p53*^*EPI*^) and *Fanca*^-/-^;*K14cre*;*Trp53*^F2-10/F2-10^ (hereinafter referred *Fanca/p53*^EPI^) animals, but not in WT or *Fanca*^-/-^ (hereinafter referred *Fanca*) ones (Table 1). SCCs were located in the lips, tongue, and oral mucosa (Fig. 1A). The incidence of SCCs was significantly higher in *Fanca/p53*^EPI^ than in *p53*^EPI^: 91% *versus* 66% (p<0.05, Fisher’s exact test) (Table 1). It is of note that double mutant *Fanca/p53*^EPI^ mice developed OSCC much earlier than *p53*^EPI^ as analyzed by a Kaplan-Meier plot (median latencies of 303 *versus* 432 days, respectively) (Fig. 1B). Tumors in the tongue and in the oral mucosa were more frequent in double mutant mice (Fig. 1A), which are the prevalent locations of HNSCC in FA patients [1, 28]. Carcinomas were well differentiated and frequently infiltrating (Fig. 2). Tumors in the lips were normally located at the mucocutaneous junction, which is a transitional region from oral mucosa to skin (Fig. 2A-C). Some animals developed more than one tumor (multicentric SCCs) (Fig. 2D-F). Therefore, Fanca and p53 cooperate to suppress SCC in the oral cavity in mice. *Fanca/p53*^EPI^ animals constitute the first animal model of spontaneous HNSCC in FA.

**Table 1.** Incidences of oral SCC in animal genotypes

| Genotype | Group size, *n* | Oral SCC, *n* (%) |
| --- | --- | --- |
| *Control* | 12 | 0 (0) |
| *Fanca* | 16 | 0 (0) |
| *p53<sup>EPI</sup>* | 12 | 8 (66) |
| *Fanca/p53<sup>EPI</sup>* | 11 | 10 (91) |

NOTE: The difference in OSCC incidence between *p53<sup>EPI</sup>* and *Fanca/p53<sup>EPI</sup>* groups was statistically significant (p-val<0.05). Statistical comparisons were performed using the Fisher's exact test.

**Figure 1.**
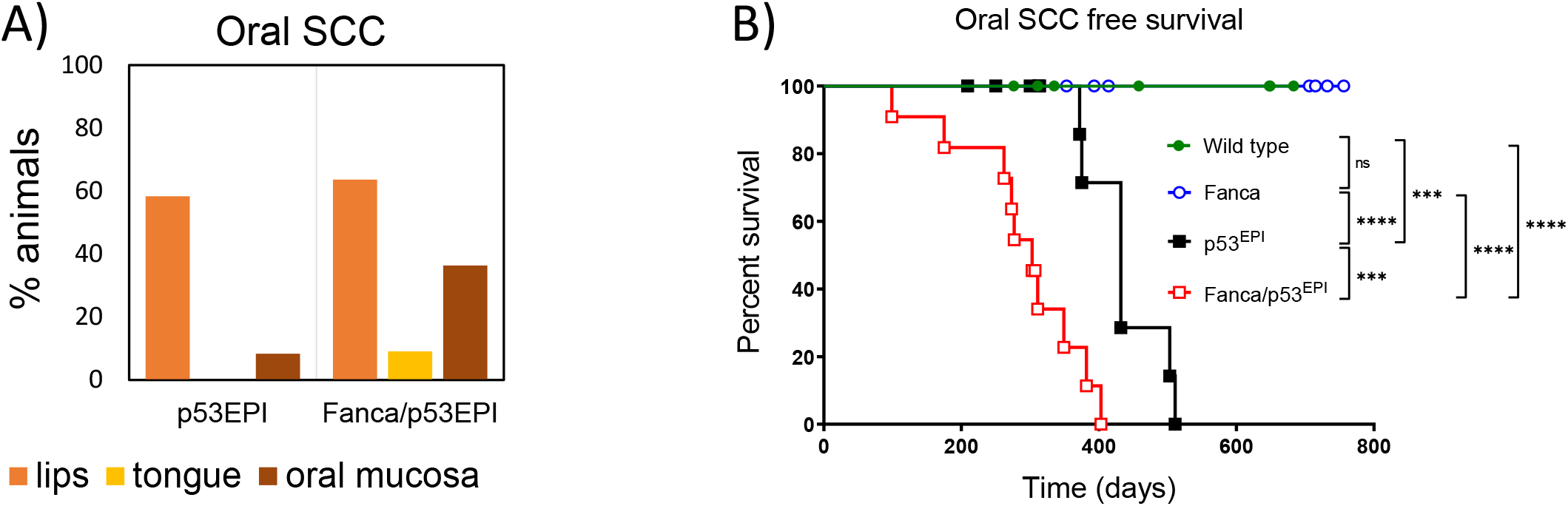
*Fanca* and *Trp53* genes cooperate to suppress oral SCC in mice. Mice from the indicated genotypes were long-term monitored (24 months), sacrificed at the observance of tumor/s development or significant morbidity, and analyzed histologically. **A)** Percentages of animals in *p53*^*EPI*^ *and Fanca/p53*^*EPI*^ genotypes having oral SCC, shown by sub-anatomical location. **B)** A Kaplan-Meier plot of the percentage of oral SCC free survival is shown. Note that *Fanca* loss accelerates SCC development in *p53*^*EPI*^ animals (*Fanca/p53*^*EPI*^ versus *Fanca*). Statistical significance between curves was determined using log rank test. ***P < 0.001, ****P < 0.0001, P > 0.05 (NS).

**Figure 2.**
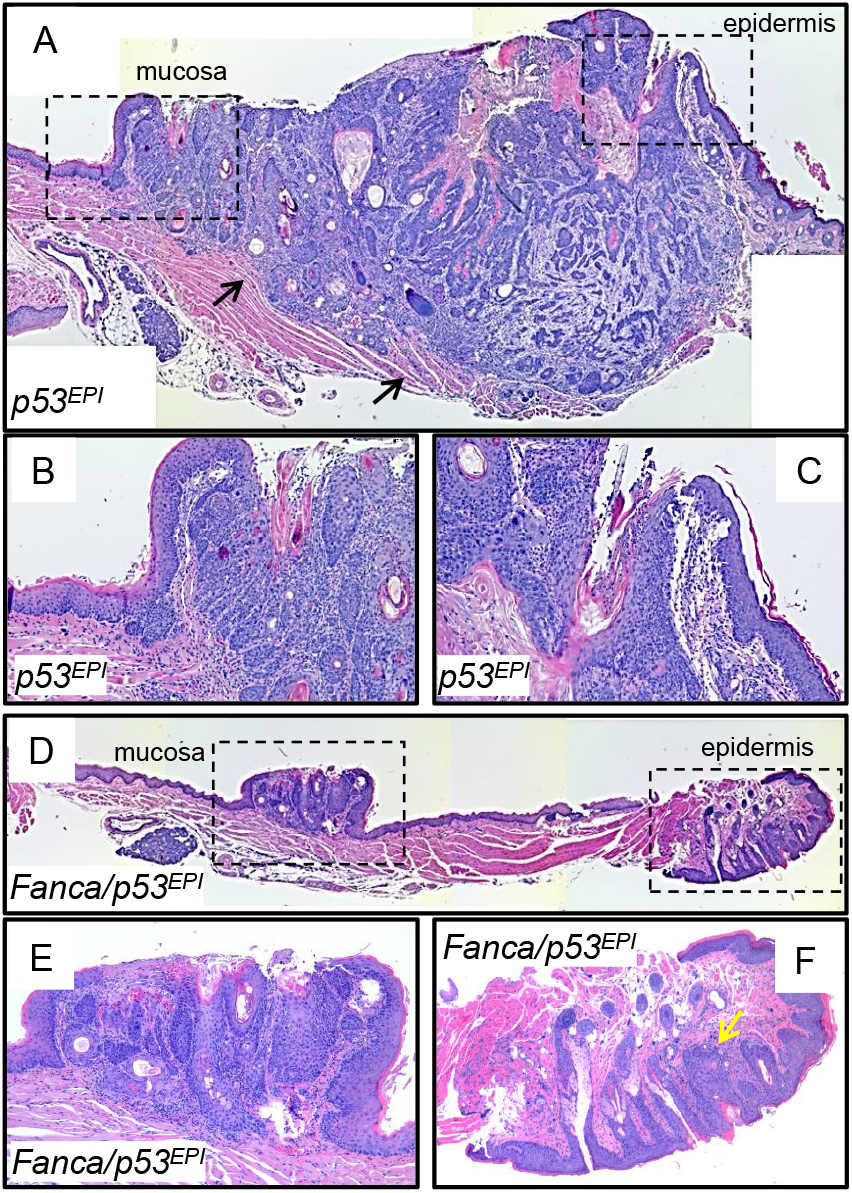
Histopathology of oral SCC in *p53*^*EPI*^ and *Fanca/p53*^*EPI*^ mice. **A)** Well differentiated SCC at the mucocutaneous junction of the lips in a *p53*^*EPI*^ mouse. The tumor infiltrates deep into the subcutaneous muscle (black arrows). Observe the mucosal (left dotted box) and epidermal (right dotted box) epithelia surrounding the carcinoma. **B)** and **C)** are magnifications of the dotted boxes in **A)**, where mucosal and cutaneous junctions are shown, respectively. **D)** Well differentiated SCCs developing at the oral mucosa and the epidermis enclosing the lips (dotted boxes) in a *Fanca/p53*^*EPI*^ animal. **E)** and **F)** are magnifications of the dotted boxes in **D)**, where mucosal and cutaneous (yellow arrow) SCCs are shown.

### OSCC from FA patients and mice express squamous differentiation markers

SCC formation involves deregulation of normal cell differentiation. As expected, immunodetection of differentiation biomarkers in tumors from FA patients (Fig. 3) showed expansion of K5-expressing cells from the basal location into suprabasal compartment in malignant cells (Fig. 3B and C). K10, a marker of suprabasal cells in epidermis that is ectopically expressed in human oral epithelial dysplasia and in skin and oral SCCs [29], was also detected in malignant cells (Fig. 3D and E). We found similar K5 and K10 staining patterns in *Fanca/p53*^EPI^ mice SCCs (Fig 4), supporting the double mutant animals as a valid model for the human disease.

**Figure 3.**
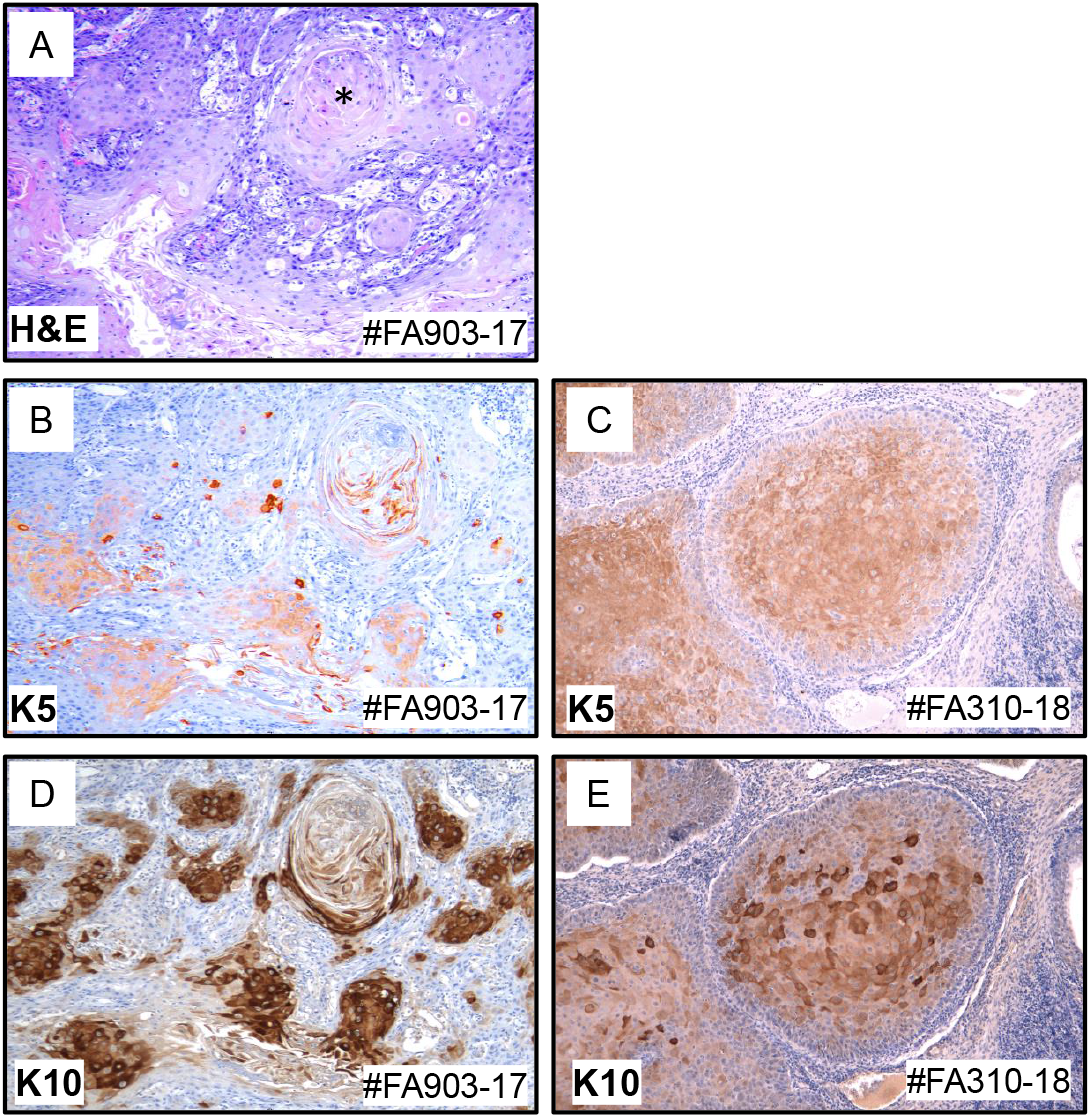
Oral SCCs in FA patients express K5 and K10 squamous differentiation keratin markers. **A)** H&E staining of a well differentiated SCC in the retromolar trigone of tumor #FA903-17, with a “horny pearl” or concentric foci of squamous differentiation (asterisk in **A**). K5 and K10 are expressed in malignant cells of the #FA903-17 SCC shown in **A)** (**B** and **D**), and in a tongue SCC (#FA310-18) from another FA patient (**C** and **E**).

**Figure 4.**
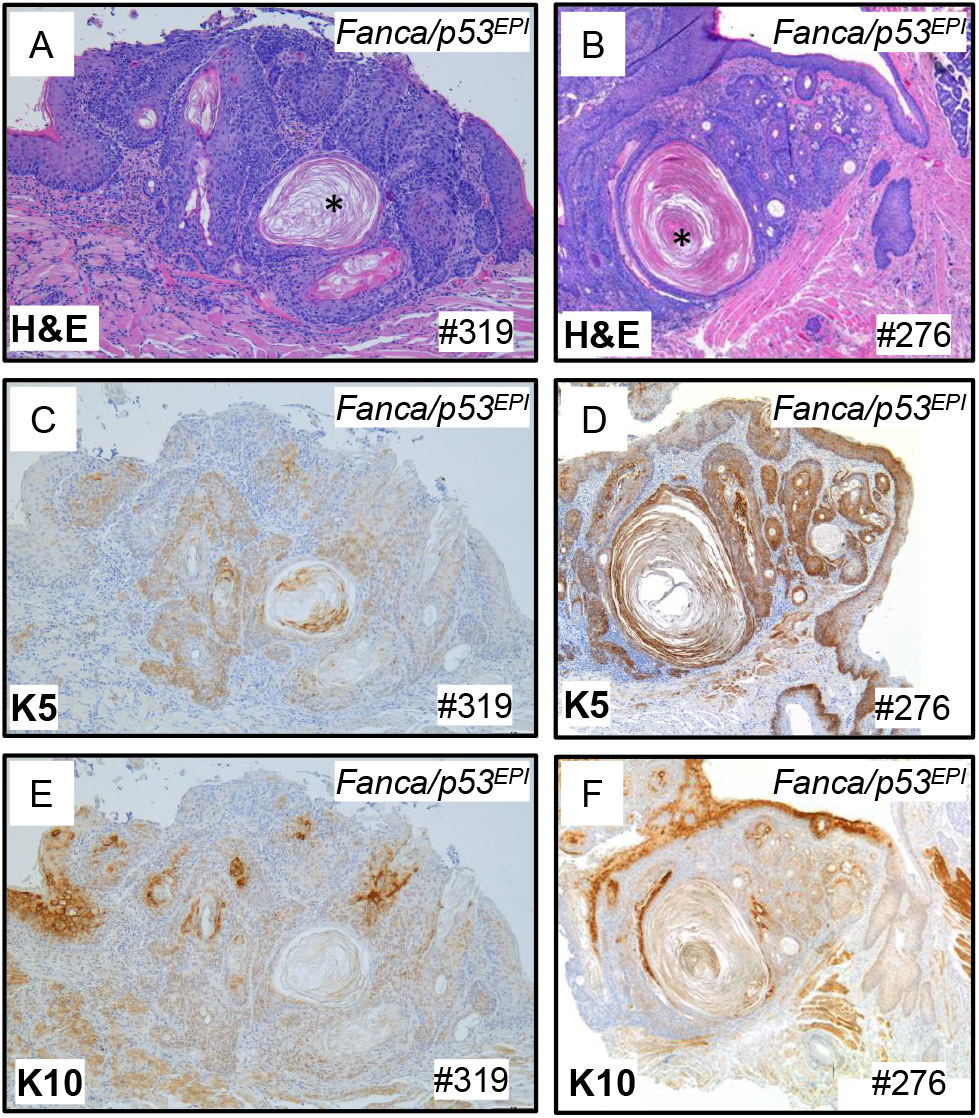
Oral SCC in *Fanca/p53*^*EPI*^ mice express K5 and K10 squamous differentiation keratin markers. **A)** and **B)** H&E staining of well differentiated SCC in the oral mucosa (A) and tongue (B) of two different *Fanca/p53*^*EPI*^ animals (#319 and #276). Both are well differentiated SCCs since “horny pearls” or concentric foci of squamous differentiation are evident in the tumors (asterisks in **A** and **B**). K5 and K10 expression is present in hyperplastic and malignant cells of the SCCs shown in **A)** (**C** and **D**) and **B)** (**E** and **F**).

### Other malignancies in *p53*^EPI^ and *Fanca/p53*^EPI^ mice

We previously reported spontaneous skin SCC development in FVB/NJ *p53*^EPI^ mice [22], with aggressive behavior and molecular signatures of poor prognostic human cancer [30]. In addition, we and others reported the development of mammary carcinomas in *p53*^EPI^ mice [31, 32]. As expected, we found skin and mammary tumors both in *p53*^*EPI*^ and in double *Fanca/p53*^*EPI*^ mice (Supplementary Fig. 1). Skin tumors were mainly SCCs, although basal cell carcinomas (BCC), sebaceous gland carcinomas and fibrosarcomas were also diagnosed. Some *p53*^EPI^ and *Fanca/p53*^EPI^ animals were sacrificed as they developed overt thymic lymphomas (Supplementary Fig. 1). These findings suggest that *Fanca/p53*^EPI^ mice also constitute a model for FA-associated cancer in locations other than head and neck.

### *Fanca* mice display frequent pre-tumoral lesions in the oral cavity

FA patients display a very high incidence of premalignant, non-invasive lesions in the oral cavity, which are difficult to prevent, diagnose and treat. We performed a careful inspection of histology samples from oral tissues collected at the time of animal sacrifice to detect possible lesions indicative of cellular transformation. Interestingly, most *Fanca* mice developed aberrant tissue phenotypes in the oral cavity (including atypia, acantolysis, picnosis or vacuolization) associated with hyperplasia or, more importantly, carcinoma *in situ* (CIS), which represents the highest atypia of premalignant epithelial cells (Supplementary Table 6 and Fig. 5). Incidences were significantly higher than in wild type (WT) animals (p-val<0.05, Fisher’s exact test), even though mean age at the analysis was very similar in both genotypes (WT=18 months *versu*s *Fanca*=19.4 months). Lesions were observed in the lips, palate, tongue, oral mucosa, and esophagus (Fig. 5 and 6), and were also found in *p53*^EPI^ and, mainly, in double *Fanca/p53*^EPI^. Some animals had more than one (multicentric) pre-tumoral lesion. Globally, the results showed that *Fanca* gene mutation is associated with development of premalignant lesions in the oral mucosa of transgenic mice.

**Figure 5.**
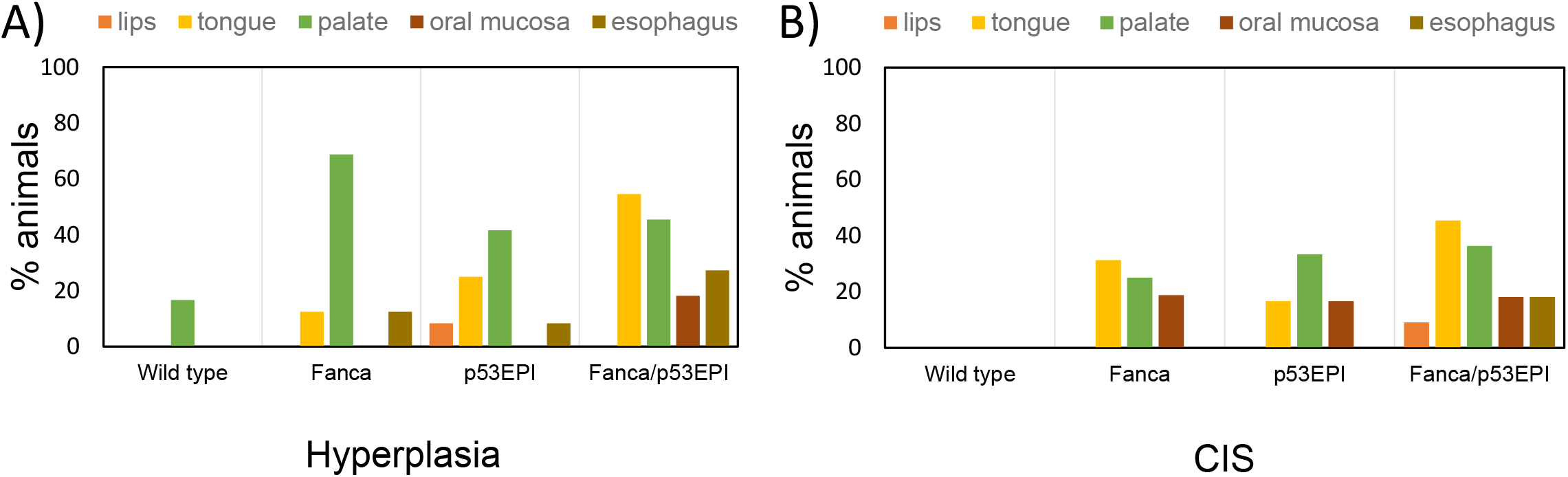
*Fanca* gene prevents the development of hyperplastic and CIS lesions in the oral cavity. Percentages of animals in all genotypes (WT, *Fanca, p53*^*EPI*^ *and Fanca/p53*^*EPI*^) showing hyperplastic **(A)** or CIS **(B)** changes in the oral cavity, shown by sub-anatomical location.

**Figure 6.**
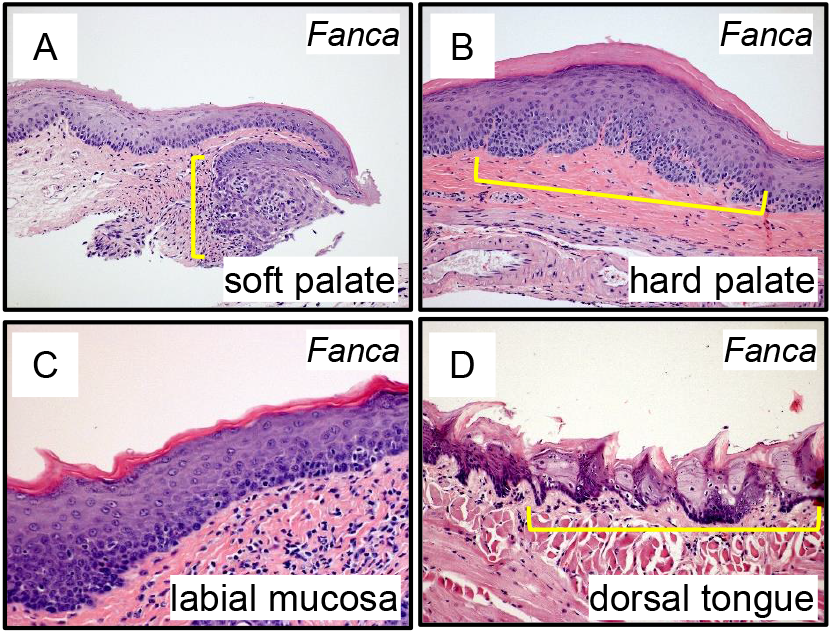
Histology of carcinoma *in-situ* (CIS) in *Fanca* mice. CIS premalignant changes found in the epithelia of the oral cavity of different *Fanca* animals. **A)** Soft palate with a focus of lichenoid inflammation, pseudocarcinomatous basal hyperplasia and severe atypia of the epithelium, characterized by nuclear pyknosis, increase of the nuclear:cytoplasm ratio, and loss of nuclear polarity of keratinocytes (square yellow bracket). **B)** Pseudocarcinomatous basal hyperplasia and severe atypia of the epithelium in the hard palate. **C)** Foci of severe atypia and epithelial hyperplasia associated to lichenoid inflammation in the labial mucosa. **D)** Foci of extreme atypia with epithelium vacuolization of the dorsal tongue filiform papillae (square bracket).

## DISCUSSION

Fanconi anemia patients display an exacerbated susceptibility to develop HNSCC early in life, mainly in the oral cavity. Due to their sensitivity to DNA damaging agents, treatments based on radiotherapy and chemotherapy should be avoided. When compared with non-FA patients, poorer survival after cancer development can be explained partially by this suboptimal treatment. Therefore, modeling the disease in animals is important to understand the oncogenic mechanisms, discover cancer drivers and biomarkers, and perform preclinical tests of new therapies. Here, we report the first mouse model of spontaneous SCC in the oral cavity with genetic depletion of a Fanconi pathway gene.

FA patients display numerous pre-malignant lesions in the oral cavity, which are difficult to diagnose and treat [2]. These lesions may lead to carcinoma development, and new methods to predict their malignant transformation are needed to improve early detection of the malignant disease [33]. We found that *Fanca* animals develop hyperplastic and CIS lesions that might resemble patient pre-malignant changes (Supplementary Table 6, Figure 5, and Figure 6). Further analysis of the lesions in *Fanca* mice and comparison with clinical samples might help to understand the mechanism of development. In addition, new chemopreventive and therapeutic approaches targeting these lesions could be tested in a preclinical setting.

Our results support the hypothesis that loss of *Fanca* gene alone does not promote the development of SCCs, even at prolonged times of observation (up to 24 months of age). Therefore, additional genetic events are needed. We have found *TP53* mutations in all five HNSCC from three FA patients (Supplementary Table 3, Supplementary Table 4), which is in agreement with the high frequency of mutations or deletions in *TP53* described in HPV-negative, FA [18-21] and non-FA patients [34-36]. In addition, *TP53* mutations are early events during HPV-negative HNSCC carcinogenesis in non-FA patients [37, 38]. These findings together with the switch from premalignant to malignant oral lesions observed in *Fanca* animals when crossed with *p53*^EPI^ (Fig. 1 and 2) point to an essential contribution of *Trp53* in early oral carcinogenesis in FA individuals. We propose that SCC development in FA individuals is a sequential process in which DNA instability produced by germline mutations in Fanconi genes might eventually give rise to alterations affecting additional cancer genes. Such double mutant cells might acquire a selective advantage over neighbor cells and expand to produce transformed clones with differential invasive capabilities.

Cooperation between *Trp53* and FA genes has already been described: tumor appearance in full body deletion in *Trp53* (*Trp53*^*-/-*^ mutant mice) is accelerated when combined with loss of *Brca2, Fancd2* or *Fancc* [14, 15, 26]. Cells from *Fanca*^*-/-*^ animals activate p53 protein function as a mechanism to control replication of cells with DNA damage [39, 40], which eventually could give rise to neoplastic cells.

Wong et al [9] reported tumor formation, mainly lymphomas, in mice with targeted disruption of exons 1 to 6 of the *Fanca* gene, which is in line with the formation of overt lymphomas in the thymus of some of our animals. However, a more systematic analysis should be performed to understand if *Fanca* or *Fanca/p53*^*EPI*^ mice have a higher incidence of such tumors when compared with WT littermates. Although the incidence and risk of skin SCC development in FA individuals have not been properly addressed, some reports showed a high incidence of BCC and SCC in FA, and at significantly younger ages than in the general population [41, 42]. As both skin SCC and BCC were observed in double *Fanca/p53*^*EPI*^ mice, we propose that these mice may also represent a model for skin SCC in FA patients. Despite a high lifetime risk of developing SCCs in the mucosae (head, neck, esophagus, vulva, or anus) in FA patients, further epidemiological studies should be performed to demonstrate that FA patients also display higher risk of development of SCCs/BCC of the skin in comparison to the general population.

In conclusion, we propose *Fanca/p53*^*EPI*^ mice as an animal model of OSCC in FA whereby future research of molecular mechanisms of the disease, discovery of novel biomarkers and preclinical testing of new therapies are warranted.

## Declaration of Competing Interest

The authors declare that they have no known competing financial interests or personal relationships that could have appeared to influence the work reported in this paper.

## Acknowledgements

The authors thank J. Bueren, P. Rio, M. Garin, and R. Gargini for helpful discussions. The authors are also indebted to the patients with FA, their families and clinicians from the Fundacion Anemia de Fanconi (Spain). The authors also thank P. Hernández and F. Sanchez-Sierra for their excellent assistance with the histological processing of the samples, and the personnel of the CIEMAT Animal Unit for the care of the mice used in this work.

## Funding

This study has been funded by Instituto de Salud Carlos III (ISCIII) through the projects PI18/00263 and P121/00208 and co-funded by FEDER and the European Union; and grants from the Spanish Fundacion Anemia de Fanconi and Fanconi Anemia Research Fund USA. The funding sources were not involved in study design; in the collection, analysis and interpretation of data; in the writing of the report; and in the decision to submit the article for publication.

## Supplementary Figure Legends

**Supplementary Figure 1**. *p53*^*EPI*^ and *Fanca/p53*^*EPI*^ mice develop overt tumors in organs such as mammary gland (**A** and **B**, adenocarcinomas), skin (**C** and **D**, SCCs; **E**, sebaceous gland carcinoma; and **F**, BCC) and thymus (**H**, lymphoma).

## Notes

### Competing Interest Statement

The authors have declared no competing interest.

