## Supplementary figures and images for "Development of a mouse model for spontaneous oral squamous cell carcinoma in Fanconi anemia"

### Supplementary Figure 1

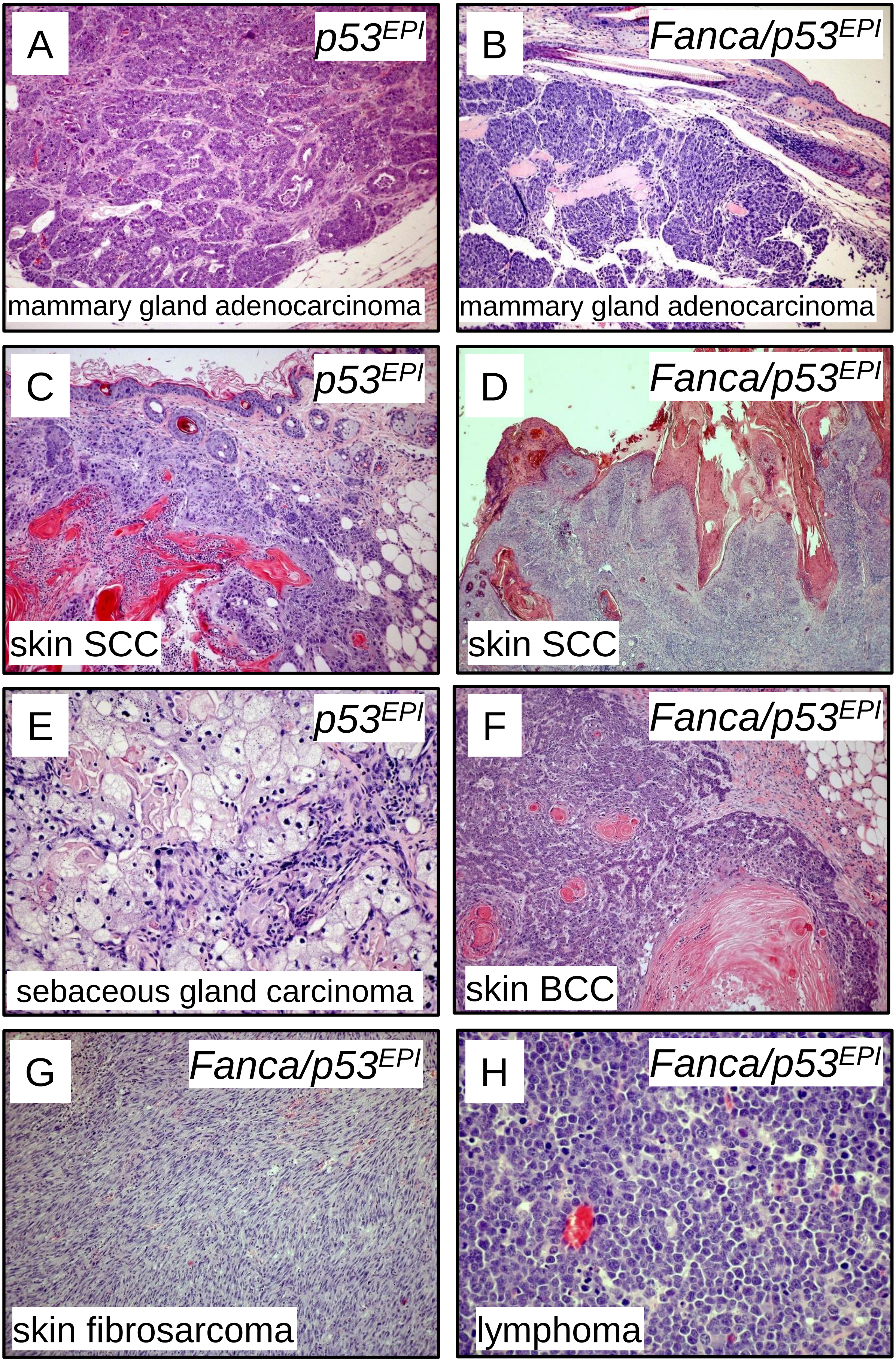
