## Supplementary Table 1 for "Development of a mouse model for spontaneous oral squamous cell carcinoma in Fanconi anemia"

**Supplementary Table 1.** Details of FA patients with OSCC

| **Patient ID** | **Age at OSCC (years)** | **Location** | **HP grading** | **Tumor ID*** | **BMT** | **Age at BMT (years)** |
| --- | --- | --- | --- | --- | --- | --- |
|  | 35 | Tongue | Well differentiated | FA310-14 |  |  |
| FA310 | 39 | Tongue | Well differentiated | FA310-18a | Yes | 8 |
|  | 39 | Tongue | Well differentiated | FA310-18b |  |  |
| FA551 | 33 | Palate | Differentiated | FA551-18 | No |  |
| FA903 | 41 | Retromolar trigone | Well differentiated | FA903-17 | No |  |

Abbreviations: HP, histopathologic; OSCC: oral squamous cell carcinoma; BMT: bone marrow transplantation.

* Tumors FA310-18a and FA310-18b are different areas from the same carcinoma.
