## Supplementary Table 3 for "Development of a mouse model for spontaneous oral squamous cell carcinoma in Fanconi anemia"

| **Sample** | **Locus** | **Gene** | **Genotype** | **AA-change** | **Type** |
| --- | --- | --- | --- | --- | --- |
| FA310-14 | chr16:89816213 | FANCA | C/A | p.Arg1055Leu | SNV |
| FA310-14 | chr16:89807256 | FANCA | AGAA/A | p.Phe1263del | INDEL |
| FA310-14 | chr17:7577581 | TP53 | A/C | p.Tyr234Asp | SNV |
| FA310-18a | chr16:89816213 | FANCA | C/A | p.Arg1055Leu | SNV |
| FA310-18a | chr16:89807256 | FANCA | AGAA/A | p.Phe1263del | INDEL |
| FA310-18a | chr17:7577570 | TP53 | C/A | p.Met237Ile | SNV |
| FA310-18b | chr16:89816213 | FANCA | C/A | p.Arg1055Leu | SNV |
| FA310-18b | chr16:89807256 | FANCA | AGAA/A | p.Phe1263del | INDEL |
| FA310-18b | chr17:7577570 | TP53 | C/A | p.Met237Ile | SNV |
| FA551-18 | chr16:89805578 | FANCA | G/C | p.Ser1377Ter | SNV |
| FA551-18 | chr16:89807256 | FANCA | AGAA/A | p.Phe1263del | INDEL |
| FA551-18 | chr17:7578190 | TP53 | T/C | p.Tyr220Cys | SNV |
| FA903-17 | chr16:89831435 | FANCA | G/A | p.Gln881Ter | SNV |
| FA903-17 | chr17:7577120 | TP53 | C/T | p.Arg273His | SNV |

**Supplementary Table 3.** FANCA and p53 mutations found in OSCC from Fanconi anemia patients.
