## Supplementary Table 6 for "Development of a mouse model for spontaneous oral squamous cell carcinoma in Fanconi anemia"

**Supplementary Table 6.** Incidence of non-invasive lesions in the oral cavity in the animal genotypes

| **Genotype** | **Group size, *n*** | **Lesions in the oral cavity, *n* (%)** | |
| --- | --- | --- | --- |
|  |  | **hyperplasia** | **carcinoma *in situ***  **(CIS)** |
| *Control* | 12 | 2 (17) | 0 (0) |
| *Fanca* | 16 | 12 (75) | 7 (44) |
| *p53^EPI^* | 12 | 7 (58) | 5 (42) |
| *Fanca/p53^EPI^* | 11 | 9 (82) | 9 (82) |

NOTE: The difference in hyperplasia or CIS incidence between *Control* and *Fanca* groups was statistically significant (p-val<0.05). Statistical comparisons were performed using the Fisher's exact test.
